# Shedding of mitochondrial Voltage-Dependent Anion Channel-1 (VDAC1) Reflects COVID-19 Severity and Reveals Macrophage Dysfunction

**DOI:** 10.1101/2025.07.07.663218

**Authors:** Marie Sjögren, Pontus Dunér, Yang De Marinis, Ariane Neumann, Karin Leandersson, Magnus Rasmussen, Claes B. Wollheim, Albert Salehi

**Author notes:** Corresponding author: Albert Salehi Department of Clinical Science Lund University, Malmö, Sweden.

## Abstract

COVID-19 severity correlates with lymphopenia and increased pro-inflammatory cytokines. However, the dysfunction of tissue macrophages in COVID-19 patients during the inflammatory cytokine storm has not been fully elucidated.

Hospitalized COVID-19 patients were divided into three groups based on their symptomatic severity: exhibiting mild, moderate, or severe symptoms. Patients exhibited successively increased serum levels of mitochondrial voltage-dependent anion channel 1 (VDAC1) at days 0, 3, 7, 10, and 14, returning to those of non-infected subjects at day 28. Serum level of VDAC1 was positively correlated with COVID-19 severity and with increased white blood cell (WBC), neutrophil, lymphocyte, procalcitonin (PCT), and gamma-glutamyltransferase (GT) levels. Peripheral blood mononuclear cells (PBMCs) from hospitalized COVID-19 patients showed increased VDAC1 content concomitant with a reduced ATP content.

Culture of monocytes, isolated from healthy individuals, and differentiated into polarized M1 macrophages, together with a cytokine mixture (IL-1β, IFN-γ, and TNF-α), to mimic the inflammatory cytokine storm, for 24 h markedly increased VDAC1 and Monocyte chemoattractant protein-1 (MCP-1) release in culture medium. The presence of the cytokine mixture reduced the ATP content, cell viability, and the phagocytic capability of macrophages. Co-staining of VDAC1 and the plasma membrane marker Na^+^/K^+^-ATPase showed that cytokine-treatment mistargeted VDAC1 to the cell surface of macrophages. All these effects were prevented by VDAC1 inhibition using VBIT-4, VDAC1-specific antibody (VDAC1-ab), or metformin.

Our findings indicate that increased VDAC1 expression and cell surface mistargeting in immune cells might be associated with cell dysfunction, potentially contributing to the severity of COVID-19 infection.

The data also indicate serum VDAC1 as a biomarker of COVID-19 severity and the use of VDAC1 inhibitors as potential drug candidates restoring macrophages and PBMCs function in individuals severely affected by COVID-19.

## Introduction

The Coronavirus disease 2019 (COVID-19) pandemic, caused by severe acute respiratory syndrome coronavirus 2 (SARS-CoV-2), mostly manifests as a mild disease. In a minority of cases, especially in the elderly or in patients with obesity and diabetes, the virus can lead to pneumonia, acute respiratory distress syndrome (ARDS), respiratory failure, and a cytokine storm with an increased risk for multiorgan failure and lethality (Huang et al. 2020; Bisen et al. 2023).

SARS-CoV-2 infection is associated with multiple changes in the activity of the peripheral immune cells, resulting in an increased inflammatory response. This increased response leads to the excessive release of pro-inflammatory cytokines, including the pleiotropic pro-inflammatory IL-1 family members interleukin-1ß (IL-1ß) and interleukin-18 (IL-18) that are generated in circulating monocytes and tissue resident macrophages, causing a cytokine storm and tissue damage. Production of IL-1ß and IL-18 is initiated through the activation of the inflammasome, a multiprotein complex, triggered by both cellular oxidative stress and the direct or indirect effects of SARS-CoV-2 (Vora, Lieberman, and Wu 2021; Dharra, Kumar Sharma, and Datta 2023). This activation results in pyroptosis, a form of inflammatory cell death (Ferreira et al. 2021; Sefik et al. 2022). Accordingly, elevated serum levels of IL-18 and Lactate Dehydrogenase (LDH) are both indicators of disease severity, with increased LDH levels reflecting cell death across various organs (Dharra, Kumar Sharma, and Datta 2023; Vora, Lieberman, and Wu 2021).

Lung tissue and monocytes from severe COVID-19 patients display increased activity of the mitochondrially bound inflammasome sensor component NLR family pyrin domain containing 3 (NLRP3) (Junqueira et al. 2022). Inhibition of NLRP3 in SARS-CoV-2-infected monocytes attenuates the production of both IL-1ß and IL-6, which are also implicated in COVID-19 pathology, but such inhibition can lead to increased viral replication (Sefik et al. 2022). Patients with type 2 diabetes (T2D) treated with metformin have reduced COVID-19 mortality (Crouse et al. 2020; Bramante et al. 2021; Xian et al. 2021; McCarthy 2023), and this has been associated with reduced NLRP3 activation (Cory et al. 2021; Singh et al. 2020). Likewise, the inhibition of NLRP3 by metformin in SARS-CoV-2-infected mice prevented lung infection (Xian et al. 2021). Thus, the proposed pharmacological tools for COVID-19 treatment (reviewed in (Duner and Salehi 2020)) often lack a focus on the underlying reasons for immune cells’ failure to target and eliminate the virus effectively.

Mitochondria not only generate ATP through oxidative phosphorylation but also play roles in calcium signalling, oxidative stress, apoptosis, and innate immunity (Smith et al. 2012). SARS-CoV-2 RNA and proteins have been shown to be localized to mitochondria, exploiting the host cell’s mitochondrial function for viral replication (Ajaz et al. 2021; Wang et al. 2022; Shteinfer-Kuzmine et al. 2024). RNA viruses are recognized by retinoic acid-inducible gene-I (RIG-I), while mitochondrial antiviral signalling (MAVS) on the outer mitochondrial membrane coordinates the innate immune response by binding RIG-I and other proteins to induce type I interferon (Wang et al. 2022; Huang et al. 2023), (reviewed in (Bhowal et al. 2023)). Conversely, MAVS is also involved in apoptosis by preventing the degradation of voltage-dependent anion channel-1 (VDAC1) (Guan et al. 2013). VDAC1 is the most abundant (reviewed in (Shoshan-Barmatz et al. 2010)) of the three outer mitochondrial membrane VDAC isoforms, facilitating the transport of metabolites (ADP and ATP), ions (calcium), and lipids. Binding of hexokinase and voltage potential regulates VDAC1 open state, determining whether metabolites or calcium are conducted through the channel (Shoshan-Barmatz et al. 2010; Shoshan-Barmatz, Shteinfer-Kuzmine, and Verma 2020; Hu et al. 2022).

Under oxidative stress and calcium overload, VDAC1 is upregulated and oligomerizes, leading to opening of the inner mitochondrial permeability transition pore (mPTP) (Tomasello et al. 2009). SARS-CoV-2 exacerbates this, generating mitochondrial reactive oxygen species (ROS) and oxidized double-stranded mitochondrial DNA (mtDNA) fragments. These fragments and other danger-associated molecular patterns (DAMPs) leak into the cytosol, activating NLRP3 and stimulator of interferon genes (STING) pathways (Hu et al. 2022; Baik et al. 2023; Xian et al. 2022; Xian and Karin 2023; Kim et al. 2019). Cyclosporin A inhibits mPTP opening, while VBIT-4 and VBIT-12 block VDAC1 oligomerization (Baik et al. 2023; Xian et al. 2022; Xian and Karin 2023; Ben-Hail et al. 2016; Verma et al. 2022). The mtDNA fragments enhance NLRP3 binding to VDAC1 oligomers (Baik et al. 2023; Xian and Karin 2023). VBIT-4 has been shown to inhibit mtDNA leakage in various cell types and disease models (Kim et al. 2019; Verma et al. 2022). RNA viruses can also induce VDAC1 oligomerization through MAVS-mediated degradation inhibition (Bhowal et al. 2023). SARS-CoV-2 alters mitochondrial-related protein (Lindeboom et al. 2024) and gene expression (Gurshaney et al. 2023), leading to reduced mitochondrial metabolism and increased glycolysis in infected nasal epithelial and immune cells, favouring viral replication (Guarnieri et al. 2023; Yeung-Luk et al. 2023). Mitochondrial ROS stabilize hypoxia-inducing factor-1α (HIF-1α), promoting glycolysis at the expense of oxidative metabolism (Guarnieri et al. 2023; Guarnieri et al. 2024; Codo et al. 2020), while HIF-1α enhances VDAC1 transcription (Guarino et al. 2020).

We previously demonstrated that mitochondrial dysfunction in type 2 diabetes (T2D) is linked to VDAC1 upregulation and mistargeting to the plasma membrane in pancreatic β-cells (Zhang et al. 2019). The impaired glucose-stimulated insulin secretion was largely due to the loss of cellular ATP through plasma membrane VDAC1 (plVDAC1). Cellular ATP and insulin secretion were restored by acute exposure to not only VBIT-4 but also to an antibody (ab) against VDAC1 as well as to metformin at 20 µM, which is within the plasma concentration range of treated patients and well below those that inhibit complex I of the respiratory chain (Emelyanova et al. 2021). Like VBIT-4, metformin rapidly reduced plasma membrane-expressed VDAC1 conductance by direct action on the channel, possibly explaining the low concentration required (Zhang et al. 2019). Of note, blockade of plVDAC1 with antibodies prevents apoptosis in neurons and endothelial cells, implicating opening of plVDAC1 in apoptosis (Akanda et al. 2008; Neumann, Kuteykin-Teplyakov, and Heumann 2024; Li et al. 2014).

In COVID-19 patients, lymphopenia and altered subsets of peripheral blood mononuclear cells (PBMCs) have been observed. A T-cell subset with elevated VDAC1 expression is prone to mitochondrial apoptosis, metabolic disruption, and senescence, correlating with lymphopenia and disease severity (Thompson et al. 2021). These dysfunctional cells are absent in healthy individuals but present in the elderly. Furthermore, immature myeloid-derived suppressor cells exhibit the highest VDAC1 expression. Treatment of PBMCs from COVID-19 patients with VBIT-4 prevented apoptosis and restored T-cell function, implicating VDAC1 in the immune cell pathology (Thompson et al. 2021). Extracellular vesicles (EVs), known to play a role in intercellular communication, have been shown to reduce SARS-CoV-2 replication and cytokine production, and notably, VDAC1 has been detected in EVs isolated from human plasma (Arteaga-Blanco et al. 2024).

Given the central role of VDAC1 in mitochondrial function, apoptosis, and immune dysregulation, we hypothesized that VDAC1 expression correlates with COVID-19 severity. In this study, we investigated VDAC1 expression in serum and PBMCs of hospitalized COVID- 19 patients and its association with disease severity and clinical parameters. We also studied human macrophages isolated from healthy individuals, exposed to inflammatory cytokines (IL- 1β, TNF-α, and INF-γ), which are induced in patients with COVID-19 (Karki et al. 2021). Expression and cellular localization of VDAC1 in the presence and absence of VDAC1 inhibitors was assessed. Macrophage functionality was evaluated by measuring phagocytic capacity, cell viability, ATP content, LDH release, and apoptosis.

## Material and methods

### Ethics

Ethical approval was obtained from the regional ethical committee at Lund University, Sweden, and all studies were conducted in accordance with the Declaration of Helsinki. Written consent from patients was obtained, ensuring data privacy and confidentiality for the use of human leukocytes from patients with COVID-19 and healthy individuals (Dnr 2020-01747, Dnr 2020- 06453). For primary cell cultures, concentrated leukocytes were obtained from blood from healthy donors (Dnr 2021-04792).

### Patient cohorts

This is a retrospective study, and the present study is a reanalysis of a material originally collected to study antibody response to SARS-CoV-2. Description of the patient group and defining criteria for the severity of COVID-19 from the first study cohort can be found in the paper by (Bläckberg et al. 2021).

The study cohort consisted of 40 hospitalized patients (>18 years of age) who were admitted to the Clinic of Infectious Diseases at Skåne University Hospital, Sweden, during 2020 and 2021 due to respiratory symptoms for COVID-19 and were positively tested for SARS-CoV-2 (confirmed by RT-PCR). Patients were divided into three groups: Group 1 (no oxygen supplementation), Group 2 (maximum of 6 L of oxygen supplementation / minute), and Group 3 (Need for more than 6 L of oxygen supplementation / minute). Peripheral blood was collected in BD Vacutainer serum tubes and set to coagulate from COVID-19 patients upon enrolment in the hospital at day 0, and at days 3, 7, 10, 14, and 28 for serum isolation. Serum was prepared by centrifugation at 150 × *g* for 10 minutes and stored at -80 °C until further use. No blood samples were obtained until the follow-up consultation on day 28 for patients discharged from the hospital during the study period.

Description of the patient group from a second study cohort and defining criteria for the severity of the COVID-19 can be found in Table 1.

**Table 1.** Demographics and clinical parameters in PBMCs from COVID-19 patient characteristics.

|  | Study Cohort | Group 1 | Group 2 | Group 3 |
| --- | --- | --- | --- | --- |
| Demographics | n = 12 | n = 5 | n = 6 | n = 1 |
| **Characteristics** |  |  |  |  |
| Age, median | 59 (31-79) | 65 (31-78) | 48 (38-72) | 79 |
| Sex, female n (%) | 5 (42) | 4 (80) | 0 (0) | 1 (20) |
| Sex, male n (%) | 7 (58) | 1 (14) | 6 (86) | 0 |
| CCI, n | 0.5 (0-3) | 2 (0-3) | 0 | 3 |
| BMI, median | 28 (24-40) | 28 (24-40) | 29 (24-37) | 26 |
| Days with COVID-19, median | 11 (6-17) | 10 (8-14) | 13 (10-17) | 6 |
| In hospital mortality (%) | 0 | 0 | 0 | 0 |
| **Clinical parameters** |  |  |  |  |
| CRP (mg/L); median | 39 (7-293) | 67 (47-140) | 22 (7-293) | 16 |
| WBC ($10^9$ c/L); median | 7 (4-16) | 6 (4-8) | 9 (4-16) | 7 |
| PMN ($10^9$ c/L); median | 5 (2-13) | 2 | 8 (2-13) | 0 |
| Lymphocytes ($10^9$ c/L); median | 1 (0-2) | 2 | 1 (0-2) | 0 |
| Platelets ($10^9$ c/L); median | 277 (133-443) | 247 (133-297) | 350 (256-443) | 0 |

Charlson Comorbidity score (CCI), Body mass index (BMI), C-reactive protein (CRP), White blood cell (WBC), Polymorphonuclear Leukocytes (PMN).
All clinical parameters are the median of the highest value for each patient (except for lymphocytes and platelets), where the lowest value for each patient is given.
Group 1: Mild disease - No oxygen, Group 2: Moderate disease - 1-6 L of oxygen/ minute, Group 3: Severe disease – More than 6 L of oxygen per minute.

Peripheral blood was collected from COVID-19 patients (n=12) in citrate tubes to isolate peripheral blood mononuclear cells (PBMCs). PBS containing 5 mM EDTA and 2.5% w/v sucrose was used to dilute concentrated leukocytes, and Ficoll-Paque Plus (GE Healthcare Biosciences, Uppsala, Sweden) gradient was used to isolate PBMCs. RIPA buffer containing protease inhibitor (Thermo Fisher Scientific, Wilmington, Delaware, USA) was used to resuspend PBMCs after centrifugation, and cells were stored at -80°C until further use. Clinical characteristics were compared between severe, moderate, and mild cases, and the VDAC1 level in the plasma was compared to plasma from non-infected healthy donors.

### Isolation of primary monocytes and macrophage differentiation

Primary monocytes were isolated from healthy blood donors as previously described (Gunnarsdottir et al. 2020). PBS containing 5 mM EDTA and 2.5% w/v sucrose was used to dilute concentrated leukocytes, and a Ficoll-Paque Plus (GE Healthcare Biosciences, Uppsala, Sweden) gradient was used to isolate PBMCs. Monocytes were isolated from PBMCs by magnetic cell sorting (MACS) using a Classical monocyte isolation kit (Miltenyi Biotec, Bergisch Gladbach, Germany) according to the manufactureŕs protocol. Thereafter, monocytes were cultured in OptiMEM, 1% Penicillin/Streptomycin (100 U/ml and 100 mg/ml) media (Gibco, Thermo Fisher Scientific, Wilmington, Delaware, USA) containing GM-CSF (10 ng/ml) for 5 days for macrophage differentiation and subsequently polarized into M1 macrophages for 2 days using GM-CSF (10 ng/ml), IFN-γ (20 ng/ml) and LPS (100 ng/ml). All cytokines used were from R&D Systems (Minneapolis, MN, USA), and LPS from Sigma-Aldrich (St. Louis, MO, USA).

### Culture of monocytes or macrophages with cytokines

Monocytes / macrophages were exposed to low cytokine level (IL-1β: 0.1 ng/ml, TNF-α 25 ng/ml, INF-γ: 25 ng/ml) or high cytokine level (IL-1β: 1 ng/ml, TNF-α: 250 ng/ml, INF-γ: 250 ng/ml); with or without VBIT-4 (20 µM), Metformin (20 µM) or VDAC1-ab (10 nM) for 24 h.

### Measurement of Cellular Viability (MTS)

Cell viability (reductive capacity) was measured using the MTS reagent kit according to the manufactureŕs instructions (Promega, Madison, WI, USA). Absorbance was measured at 490 nm using a microplate reader (CLARIOstar Plus, BMG LABTECH, Ortenberg, Germany). Cell viability was expressed as a percentage of the absorbance in the treated groups compared to the control group.

### MCP-1 and VDAC-1 measurement

Human CCL2/MCP-1 ELISA (R&D systems, Minneapolis, MN, USA) was used to quantify released MCP-1 from monocytes and macrophages. Analyses were performed according to instructions from the manufacturer. Due to low levels of released MCP-1 in the absence of an inflammatory stimulus, an undetectable level was considered as 0. VDAC1 protein was determined in serum and cell homogenates or cell culture medium with Human VDAC1/PORIN ELISA (LSBio, Shirley, MA, USA) according to the manufactureŕs instructions. The quantity of protein in cell culture experiments was determined by Pierce BCA Protein Assay Kit (Thermo Fisher Scientific, Wilmington, Delaware, USA) according to the manufactureŕs protocol. VDAC1 and MCP-1 values in the cell medium were normalized to the amount of cell protein in each sample.

### Immunofluorescence analysis by confocal microscopy

Monocytes or macrophages were cultured in gelatine-coated (Thermo Fisher Scientific, Wilmington, Delaware, USA) Nunc™ Lab-Tek™ II Chamber Slide™ System (Thermo Fisher Scientific, Wilmington, Delaware, USA) in the presence or absence of indicated agents. Thereafter, they were washed twice with cold PBS and then fixed with ice-cold 100% Methanol for 5 min at room temperature (RT). The fixed cells were washed in cold PBS and then blocked in 5% fetal bovine serum in PBS for 1 h in RT. Primary antibodies against rabbit monoclonal VDAC1 (Abcam; Ab154856; 1:200) and mouse monoclonal alpha 1 Sodium Potassium ATPase antibody [464.6] (Abcam; Ab7671; 1:200) diluted in blocking buffer were applied and incubated overnight at +4°C. After washing several times with PBS, cells were incubated for 1 h in blocking solution containing 0.02% Triton X-100, secondary Donkey AF488 anti-mouse (Jackson ImmunoResearch; 715-545-150; 1:200) and Donkey Cy3 anti-rabbit (Jackson ImmunoResearch; 711-165-152; 1:200) antibody and Hoechst 33258 (1:1000). The cells were washed with PBS, coverslips were mounted with mounting media, and visualized by confocal microscopy (Carl Zeiss, Germany) with a 40x magnification. The intensity of VDAC1 and Hoechst was visualized by confocal microscopy (Carl Zeiss, Germany), and colocalization analysis of VDAC1 and Na^+^/K^+^ ATPase was quantified using software ZEN 2.3 blue edition.

### Cell apoptosis analysis

Cell apoptosis was examined spectrofluorometric by Hoechst 33258 staining as described elsewhere (Majtnerova et al. 2021). After exposure of macrophages to different experimental conditions, the cells were fixed and stained as described (Majtnerova et al. 2021)Hoechst intensity was visualized by confocal microscopy (Carl Zeiss, Germany) and analysed using ZEN 2.3 blue edition software. Apoptosis was expressed as a percentage of the mean Hoechst intensity in all groups compared to the control group.

### ATP determination

ATP content was determined in isolated PBMCs from COVID-19 patients and healthy controls, and in monocyte-derived macrophages exposed to inflammatory cytokines (IL-β, TNF-α, and IFN-γ) for 24h in the absence or presence of VBIT-4 (20 µM), Metformin (20 µM), or VDAC1- ab (10 nM) using a luminescence assay kit according to the manufacturer’s protocol (Promega, Madison, WI, USA). Protein levels in the samples were measured by Pierce BCA Protein Assay Kit (Thermo Fisher Scientific, Wilmington, Delaware, USA) according to the manufactureŕs protocol. ATP values were normalized to the amount of cell protein in each sample.

### LDH release assay

LDH release was measured in monocyte-derived macrophages exposed to inflammatory cytokines (IL-β, TNF-α, IFN-γ) for 24 h in the absence or presence of VBIT-4 (20 µM), Metformin (20 µM), or VDAC1-ab (10 nM), according to the manufacturer’s protocol (Promega, Madison, WI, USA). LDH values were normalized to the amount of cell protein (measured by Pierce BCA Protein Assay Kit (Thermo Fisher Scientific, Wilmington, Delaware, USA) in each sample and were expressed as a percentage in the treated groups compared to the control group.

### Detection of phagocytosis by FITC-dextran uptake analysis

To measure the phagocytic ability in macrophages, the FITC-dextran uptake assay was set up by incubating the cells with FITC-dextran (Sigma-Aldrich, St. Louis, MO, USA) in duplicate. Purified monocytes were cultured in 96-well plates with a concentration of 0.1 x 10^5^ cells/well, differentiated into M1 polarized macrophages, and exposed to cytokines (IL-1β: 1 ng/ml, TNF- α: 250 ng/ml, INF-γ: 250 ng/ml) with or without VDAC1 inhibitors described above. FITC-dextran (0.25 mg/ml) was added to each well and incubated at +37°C for 30 min. After incubation, the cells were washed extensively using cold PBS to remove excess FITC-dextran.

Macrophages were detached using trypsin. Medium containing FBS was added to neutralize the trypsin, and the cells were spun down at 400 x g for 2 min at +4°C. All liquid was removed and replaced with FACS buffer. FACS analysis was performed using a CytoFlex Flow cytometer (Beckman Coulter, USA), and median fluorescence intensity (MFI) was measured. MFI was expressed as a percentage in the treated groups compared to the control group.

### Statistical analysis

Statistical analyses were performed using GraphPad Prism (GraphPad Software Inc., version 9, San Diego, CA, USA) or IBM SPSS Statistics, version 24.0 (IBM Corp., Armonk, NY). The results are expressed as means ± SEM for the indicated number of observations or illustrated by a representative observation of the result obtained from different experiments (confocal microscopy). Differences between two groups were analysed using Student’s t-test, and for multiple groups, one-way ANOVA followed by Tukey’s multiple comparisons test was used. P-value <0.05 was considered significant.

## Results

### Clinical characteristics in COVID-19 patients

The present study reanalyses a material collected initially to study the antibody response to SARS-CoV-2. Description of the patient group and defining criteria for the severity of COVID- 19 from the first study cohort can be found in the paper by (Bläckberg et al. 2021)).

Forty COVID-19-infected and hospitalized patients admitted to the Clinic of Infectious Diseases, Lund, Sweden, were included in the study. Based on the severity of the disease, the patients were divided into three groups; group 1 patients exhibiting mild symptoms with no oxygen treatment (n=18), group 2 patients with moderate COVID-19 requiring oxygen (1-6L / min) (n=16), and group 3 patients with severe COVID-19 (n=6) involving non-invasive ventilation, high-flow nasal cannula (HFNC) oxygen therapy, ICU treatment and / or death. Serial blood samplings were obtained on day 0 (n = 40), 3 (n = 25), 7 (n = 11), 10 (n = 6), 14 (n = 5), and day 28 (n = 36). The group with severe COVID-19 symptoms showed higher C-reactive protein (CRP), PCT, and IL-6 levels than the non-severe group. Fatal and life-threatening symptoms were only observed in group 3. Two patients died during the study period before follow-up blood sampling on day 28 (Bläckberg et al. 2021).

An additional cohort (n=12) of COVID-19-infected and hospitalized patients was included, and basic characteristics and clinicopathological features were analysed using the same grouping strategy based on the severity of the disease (Table 1).

In this cohort, only one patient was included in group 3 requiring HFNC oxygen therapy, without the need for intensive care. Blood samples were taken on day 0, and different clinical parameters, e.g., CRP, procalcitonin, GT, WBC, lymphocytes, and neutrophils, were measured. The median age for this smaller cohort was 59 years (IQR 31-79), in which the majority were men (58%). Patients in group 1, experiencing mild COVID-19 symptoms, were 80% women and 14% men, while group 2, with moderate symptoms, consisted of 86% men and no women. One patient from group 2 experienced pulmonary embolism. The median number of days with COVID-19 for all patients was 11 (IQR 6–17). No patients in this cohort died.

### VDAC1 correlates with severity and clinical parameters in COVID-19 patients

Since a study (by (Thompson et al. 2021)). showed an increased VDAC1 expression in T-cells, accompanied by gene programs and functional characteristics linked to mitochondrial dysfunction and apoptosis in COVID-19 patients, we analysed VDAC1protein in serum from the first cohort (Bläckberg et al. 2021), and PBMCs isolated from COVID-19 patients in the second cohort (Table 1). As shown in Figure 1, VDAC1 level in the serum was increased during the progression of the COVID-19 infection (Figure 1A), with its highest peak at day 10. On day 28, VDAC1 levels in recovered COVID-19 patients were decreased to the basal level seen in non-infected controls (Figure 1A). The magnitude of VDAC1 levels also depended on the severity of the COVID-19 infection (Figure 1B). Thus, the highest serum level of VDAC1 was observed in group 3, representing patients with severe COVID-19 symptoms (Figure 1B). Next, we performed a correlation analysis between VDAC1 expression in serum and clinical parameters of COVID-19 patients. We observed a positive correlation between VDAC1 levels in serum and WBC (p<0.0001), neutrophils (p=0.014), lymphocytes (p=0.0003), GT (p=0.017), and PCT (p=0.00045) (Figure 1C-G).

**Figure 1.**
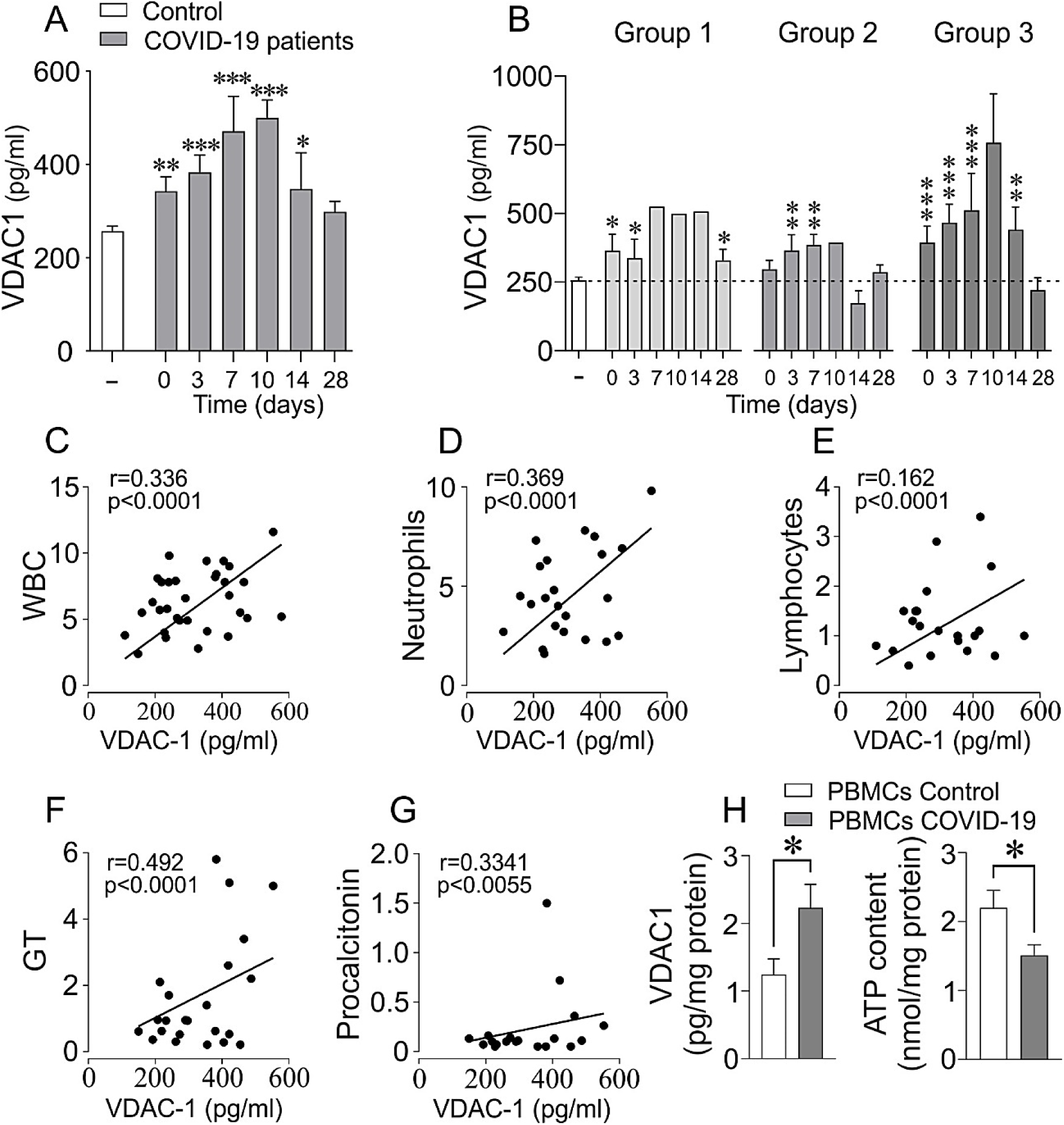
Increased VDAC1 level in serum correlates with disease progression and severity. Peripheral blood was collected from hospitalized COVID-19 patients on day 0 (n = 40), 3 (n = 25), 7 (n = 11), 10 (n = 6), 14 (n = 5), and day 28 (n = 36) for isolation of serum. Serum VDAC1 level was measured in all COVID-19 patients and compared with healthy individuals, showing increased levels during the progression of the disease **(A)**. The COVID-19 patients were divided into three groups depending on the severity of the disease: Group 1: mild disease, no oxygen needed; Group 2: moderate disease, 1-6L oxygen needed; Group 3: severe disease, intensive care was needed. The magnitude of the VDAC1 level was dependent on the COVID-19 disease severity **(B)**. Clinical parameters were compared to VDAC1 level in serum, and simple linear regression shows the correlation between VDAC1 and WBC **(C)**, neutrophils **(D)**, lymphocytes **(E)**, GT **(F)**, PCT **(G)**. Peripheral blood was collected from the second cohort of hospitalized COVID-19 patients (n=12) for isolation of PBMCs. VDAC1 levels and ATP content **(H)** were measured and compared to healthy individuals (n=8). Data represent mean ± SEM. *p < 0.05, **p < 0.01, ***p < 0.001.

VDAC1 protein levels and ATP content were also analysed in PBMCs isolated from COVID- 19 patients and compared to non-infected controls by ELISA. We found a 79% increase in cellular content of VDAC1 levels (p=0.04) concomitant with a 46% decrease in cellular ATP content in COVID-19 patients (p=0.02) (Figure 1H). These results demonstrate that increased VDAC1 expression in PBMCs correlates with disease progression and severity in patients with COVID-19.

### Cytokine-induced increased VDAC1 expression and its translocation to the plasma membrane in macrophages are prevented by VDAC1 inhibitors

Since poorly regulated blood sugar is seen in diabetic subjects, it is an independent risk factor for COVID-19-related mortality (Crouse et al. 2020), we next evaluated the impact of a high glucose concentration on cellular viability in polarized M1 macrophages cultured for 72 h. Our data showed that the ambient glucose level of culture medium is an essential factor for macrophage functionality, since culture at 20 mM glucose reduced cellular viability of macrophages compared to 5 mM glucose (Figure S1A). Therefore, in the following experiments, we used a culture medium containing 5 mM glucose to investigate the possible effect of the high cytokine-mixture on the subcellular localization of VDAC1 in macrophages by confocal microscopy in the presence or absence of VBIT-4 during cytokine challenge.

Given the significantly elevated cytokine levels in tissue observed in COVID-19 patients (Huang et al. 2020; Chen et al. 2020; Zhu et al. 2020; Del Valle et al. 2020; Ye, Wang, and Mao 2020), we next cultured polarized M1 macrophages for 24 h to test the effect of two different concentrations of cytokine-mixture on the VDAC1 protein level in relation to cell viability. The cytokines chosen were IL-1β, TNF-α, and INF-γ, which has all been shown to be induced in COVID-19 patients (Faraj and Jalal 2023; Karki et al. 2021) and also known to be involved in promoting inflammatory macrophage polarization in COVID-19 (Zhang et al. 2020). As seen in Figure S1B, while the presence of low cytokine-mixture (IL-1β: 0.1 ng/ml, TNF-α 25 ng/ml, INF-γ: 25. ng/ml) did not affect VDAC1 protein levels, a 10-fold higher concentration of cytokine-mixture (IL-1β: 1 ng/ml, TNF-α: 250 ng/ml, INF-γ: 250 ng/ml) significantly increased VDAC1 protein levels in monocyte-derived macrophages. Additionally, the increase in VDAC1 protein expression was markedly prevented by the presence of VBIT-4 during culture period (Figure S1B). Evaluation of cell viability showed that higher concentration of cytokine-mixture indeed reduced the cellular viability of macrophages, which was significantly prevented by the presence of VBIT-4 (Figure S1C). Considering our experimental findings that low cytokine levels during the culture period did neither significantly affect VDAC1 levels nor cell viability in macrophages (Supplementary Figure 1B and 1C), we opted to investigate the impact of high-dose cytokines on macrophage functionality, specifically focusing on VDAC1 expression and cellular localization. As seen in Figure 2, VDAC1 expression (green) was almost entirely confined to the cytoplasm without any significant membrane localization in macrophages cultured at 5 mM glucose. Culture of macrophages with a high concentration of inflammatory cytokines brought about a marked mistargeting of VDAC1 to the cell surface, as indicated by its co-localization with the membrane marker (Na^+^/K^+^ ATPase; red) (Figure 2 and Figure S2A). The VDAC1 inhibitor, VBIT-4 (20 µM) prevented this effect of cytokines on VDAC1 expression and its mistargeting to the cell surface (Figure 2, Figure S2A). Culture with cytokine-mixture also increased nuclear pro-apoptotic changes in macrophages as is shown by measuring the intensity of nuclear Hoechst 33258 staining (Figure 2B and Figure S2B). The apoptotic effect of cytokines on macrophages was prevented by VBIT-4 (20 µM) (Figure 2 and Figure S2B).

**Figure 2.**
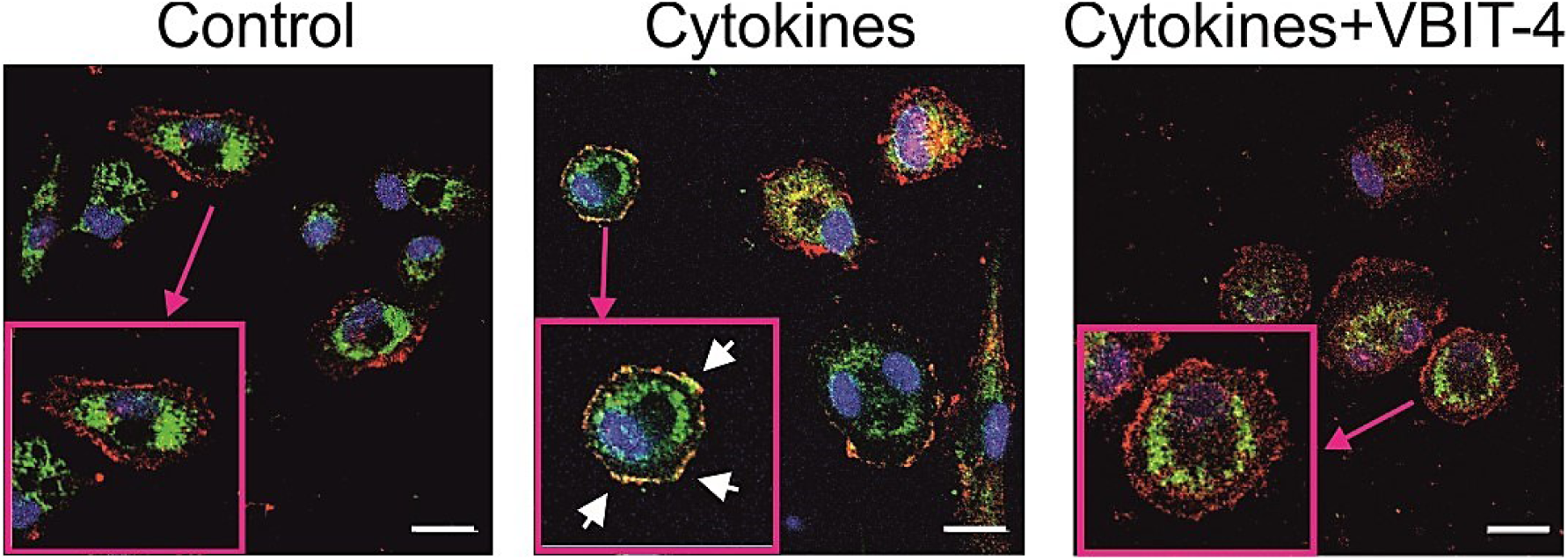
VBIT-4 prevents translocation to the plasma membrane in macrophages. Representative confocal immunofluorescence images from 3 independent experiments of VDAC1 (green) expression and its co-staining with the membrane marker Na^+^/K^+^ ATPase (red), and Hoechst (blue) Possible effects of cytokine mixture ± VBIT-4 (20 µM) in macrophages on the cellular localization of VDAC1 was examined by confocal microscopy after 24 h of the culture period. The culture of macrophages exposed to the cytokine mixture not only increased VDAC1 expression but also resulted in a marked mistargeting of VDAC1 to the cell surface, as indicated by its co-localization with the membrane marker Na^+^/K^+^ ATPase. The illustration shows an increased size of the cells. Scale bars indicate 20 µm.

### VDAC1 inhibitors restore cellular function in monocytes and macrophages

VDAC1 and MCP-1 release was measured in medium from monocytes and macrophages cultured with the high cytokine-mixture (IL-1β: 1 ng/ml, TNF-α: 250 ng/ml, INF-γ: 250 ng/ml) for 24 h in the presence or absence of VBIT-4. Cytokine-mixture significantly increased VDAC1 release in both monocytes (Figure S3A) and macrophages (Figure 3A), which was prevented by VBIT-4 (Figure S3A and Figure 3A). MCP-1 release increased significantly only from macrophages (Figure 3B), whereas there was a tendency of increased MCP-1 release from monocytes (Figure S3B), although it did not reach a significant level (p<0.06). To study the possible consequence of VDAC1 mistargeting to the cell surface in macrophages, we measured ATP content, cell viability, LDH release and phagocytic capacity after 24 h exposure to the cytokine-mixture in the presence or absence of VBIT-4 (20 µM), Metformin (20 µM), or VDAC1-ab (10 nM), and compared to basal culture condition (5 mM glucose) (Figure 3C-F). Macrophages exposed to the cytokine-mixture were found to have 47.2% decrease in ATP content (p<0.001), indicative of ATP loss via mistargeted VDAC1 to the plasma membrane since the ATP loss was prevented by VDAC1 inhibitors (VBIT-4, 20 µM) (p<0.05), Metformin (p<0.01), or VDAC1-ab (p<0.01) (Figure 3C)). Macrophages exposed to the cytokine-mixture also displayed a 55.7% decrease in cell viability (Figure 3D), which was also prevented and restored to control level when VBIT-4 (p<0.001), Metformin (p<0.001), or VDAC1-ab (p<0.001) were present during the culture period.

**Figure 3.**
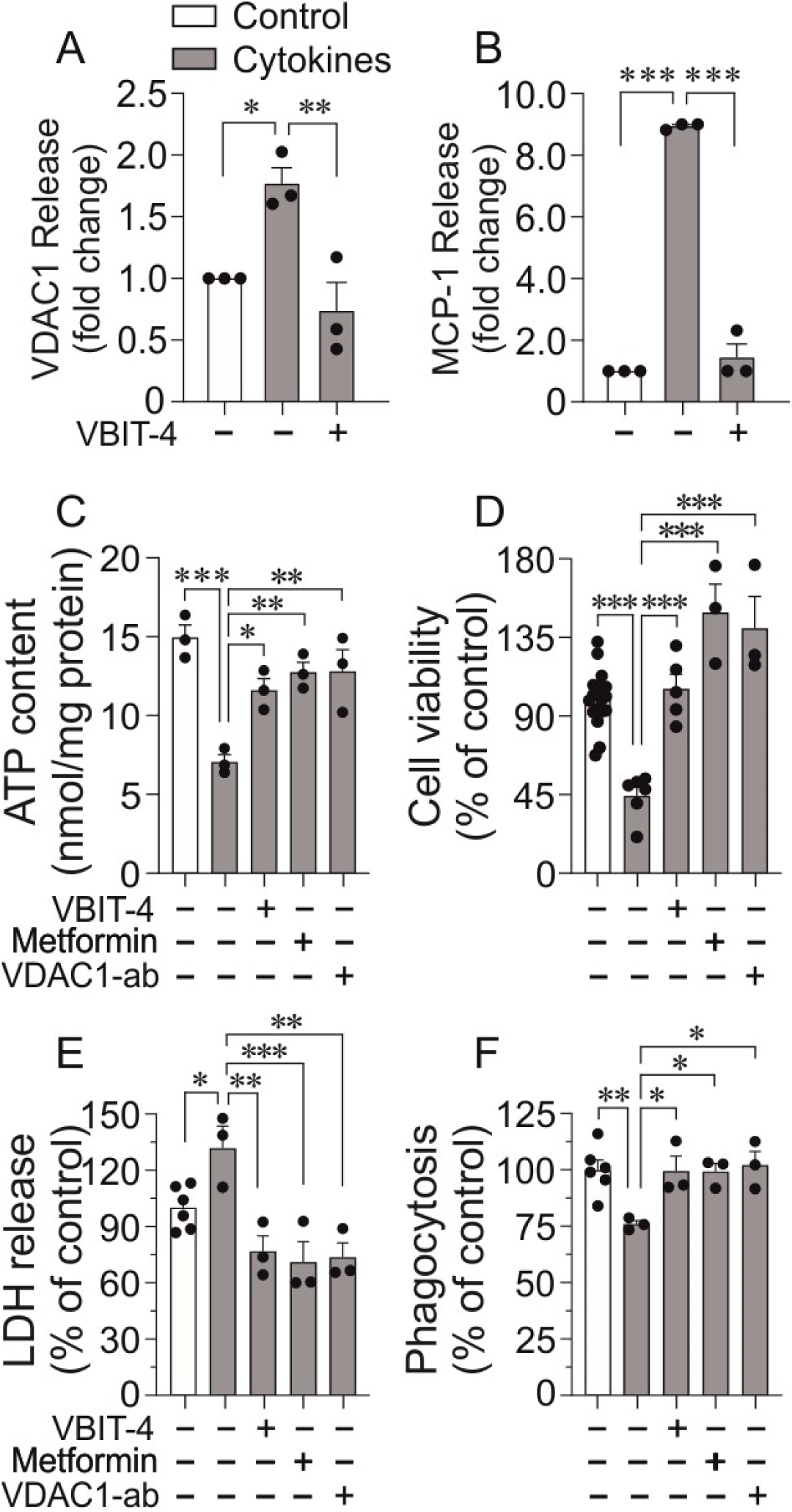
VDAC1 inhibitors restore cellular function in macrophages. The impact of cytokines on macrophage function in the presence or absence of VDAC1 inhibition. Shown is the effect of cytokine mixture (IL-1β: 1 ng/ml, TNF-α: 250 ng/ml, INF-γ: 250 ng/ml) on VDAC1 and MCP-1 release in the presence or absence of VBIT-4 (20 µM) and compared to basal conditions (5 mM glucose). VBIT-4 significantly prevented the increased VDAC1 and MCP-1 release **(A, B)**. The effect of cytokine mixture on ATP content in macrophages cultured for 24 h in the presence or absence of VBIT-4 (20 µM), metformin (20 µM), or VDAC1 antibody (10 nM). For comparison, ATP content at basal conditions (5 mM glucose) is included **(C)**. Cell viability measured as reductive capacity by MTS in macrophages cultured for 24 h with cytokine mixture in the presence or absence of VBIT-4 (20 µM), metformin (20 µM), or VDAC1 antibody (VDAC1-ab) (10 nM) compared to basal conditions (5 mM glucose) **(D)**. Lactate Dehydrogenase (LDH) release from macrophages cultured with the cytokine mixture for 24 h in the presence or absence of VBIT-4 (20 µM), metformin (20 µM), or VDAC1 antibody (10 nM) (**E**). Phagocytic capacity measured by flow cytometry in macrophages cultured with cytokine mixture for 24 h in the presence or absence of VBIT-4 (20 µM), metformin (20 µM), or VDAC1 antibody (10 nM) **(F)**. Data represent mean ± SEM from 3-4 independent experiments. *p<0.05, **p < 0.01, ***p<0.001.

Culture of monocytes with the cytokine-mixture for 24 h showed a decreased cell viability, measured by the MTS assay, at the high concentration (IL-1β: 1 ng/ml, TNF-α: 250 ng/ml, INF- γ: 250 ng/ml) as compared to control or low concentration of cytokines (IL-1β: 0.1 ng/ml, TNF- α: 25 ng/ml, INF-γ: 25 ng/ml) group (Figure S3C). The presence of VDAC1 blocker (VBIT-4) (20 µM) attenuated this effect of high cytokine-mixture and restored cellular viability to control level (Figure S3C).

To further assess the impact of the high cytokine concentration mixture (IL-1β: 1 ng/ml, TNF- α: 250 ng/ml, INF-γ: 250 ng/ml) on toxic and apoptotic effects in macrophages, we analysed the release of lactate dehydrogenase (LDH) into the culture medium after exposure of macrophages to the cytokines for 24 h. LDH is rapidly released into the cell culture medium upon plasma membrane disruption and is a widely used marker for cytotoxicity and cell death (Kumar, Nagarajan, and Uchil 2018). Macrophages with the cytokine-mixture released more LDH (p<0.05) compared to those cultured at basal conditions (Figure 3E). This effect of cytokine-mixture was prevented by VBIT-4 (p<0.01), Metformin (p<0.001), VDAC1-ab (p<0.01) (Figure 3E).

Macrophages, along with dendritic cells and neutrophils, are professional phagocytes forming the first line of defence against invading pathogens (Campbell and Colgan 2015; Parkos 2016; Shalova et al. 2015). A decreased phagocytic capacity is found in patients with COVID-19 that is associated with the severity of the disease (Nomani et al. 2021; Peyneau et al. 2022). Phagocytic capacity, measured by FITC-dextran uptake analysis, showed a 24% decrease (p<0.01) in macrophages exposed to the high cytokine-mixture (IL-1β: 1 ng/ml, TNF-α: 250 ng/ml, INF-γ: 250 ng/ml) (Figure 3F). The reduction of phagocytic capability induced by cytokine-mixture was prevented and restored to the control level by VBIT-4 (p<0.01), Metformin (p<0.01), and VDAC1-ab (p<0.01) (Figure 3F).

## Discussion

The results of our study align with the growing evidence that mitochondrial dysfunction, particularly involving VDAC1, plays a critical role in the pathogenesis of severe COVID-19 disease. Elevated VDAC1 expression was observed in the serum and PBMCs from COVID-19 patients, with the highest levels seen in those experiencing severe symptoms. These findings reflect the broader context of immune dysregulation and cellular stress that defines the pathophysiology of severe COVID-19 cases, reinforcing the notion that VDAC1 is not only a marker of disease severity but potentially a driver of the immune dysfunction and inflammation seen in these patients.

One of the key observations in this study is the correlation between VDAC1 levels and clinical parameters such as WBC, neutrophils, lymphocytes, PCT, and GT. The strong positive correlation between VDAC1 levels and these markers of inflammation and immune response underscores the potential role of increased VDAC1 in the excessive inflammatory response characteristic of severe COVID-19. This is consistent with previous findings that VDAC1 dysregulation is linked to mitochondrial damage and immune cell apoptosis in COVID-19 patients (Thompson et al. 2021). Our findings extend this understanding by demonstrating that VDAC1 levels correlate with clinical severity, suggesting that VDAC1 could serve as a prognostic biomarker for identifying patients at higher risk for severe disease.

The mechanism by which VDAC1 contributes to disease progression appears to be multifaceted. VDAC1, a key regulator of mitochondrial metabolism and apoptosis, is upregulated under oxidative stress and inflammation conditions, which are hallmark features of severe COVID-19 (Ajaz et al. 2021; Ferreira et al. 2021). In our study, increased VDAC1 expression in PBMCs was accompanied by a significant reduction in cellular ATP content, suggesting a role for VDAC1 in ATP loss and energy dysregulation. This ATP depletion may be related to the mistargeting of VDAC1 to the plasma membrane, where it forms channels that disrupt cellular ATP homeostasis, as previously reported for β-cells dysfunction in human and animal models of T2D (Zhang et al. 2019) and also in retinal cells dysfunction causing diabetic retinopathy (Tariq, Sjögren, and Salehi 2024). Our data showing ATP loss in PBMCs from COVID-19 patients supports the idea that mitochondrial dysfunction, mediated by the increased VDAC1 expression and its cell surface mistargeting, is central to the cellular damage observed in severe cases of the disease. Our present findings are particularly relevant to the recent and ongoing pursuit of VDAC1 as a target for the prevention of T2D (Zhang et al. 2019) and its devastating complication, i.e., retinopathy (Tariq, Sjögren, and Salehi 2024).

In addition to cellular energy dysregulation, we observed that VDAC1 mistargeting to the plasma membrane in macrophages exposed to inflammatory cytokines resulted in significant cellular dysfunction. Cytokine-induced VDAC1 translocation was associated with increased nuclear pro-apoptotic changes, reduced cell viability, decreased ATP content, and impaired phagocytic capacity. This suggests that increased expression and altered localization of VDAC1 not only contribute to pro-apoptotic changes but may also be associated with the functional impairment of macrophages, which are essential for controlling viral infections. In severe COVID-19, the failure of macrophages to neutralize and destroy the virus effectively could exacerbate viral replication and inflammation, leading to the cytokine storm and multi-organ damage commonly seen in critical patients (Nomani et al. 2021; Shalova et al. 2015). Increased cellular VDAC1 expression induced by pathological conditions is associated with its oligomerization, leading to mitochondrial DNA (mtDNA) leakage (Xian and Karin 2023). A recent study has shown that lymphocytes and a specific subset of T-cells detected in COVID- 19 patients exhibit mitochondrial dysfunction and are susceptible to apoptosis due to elevated VDAC1 levels (Thompson et al. 2021). Interestingly, T-cell apoptosis is prevented in vitro by VDAC1 blocker VBIT-4 (Thompson et al. 2021). An increased VDAC1 protein expression is associated with its mistargeting to the cell surface in non-immune cells (Tariq, Sjögren, and Salehi 2024; Zhang et al. 2019; Mohammad Al-Amily et al. 2023). Our data thus suggest that VDAC1 overexpression would lead to the formation of a mega channel through its oligomerization that reportedly causes mtDNA release (Kim et al. 2019; Smilansky et al. 2015). Thus, mtDNA leakage membrane translocated VDAC1 into systemic blood could indeed exacerbate and further amplify immune responses that would exacerbate the already excessive pro-inflammatory cytokine release in COVID-19 patients. Buttressing this assertion, a very recent study by Shteinfer-Kuzmine et al. has shown that COVID-19 patients display a markedly increased serum level of mtDNA (Shteinfer-Kuzmine et al. 2024). They also showed that the envelope protein of SARS-CoV-19 increases VDAC1 protein and causes its oligomerization, leading to mtDNA leakage into the cytosol of human neuroblastoma cells, which was inhibited by blocking VDAC1 oligomerization with VBIT-4 (Shteinfer-Kuzmine et al. 2024).

The use of VDAC1 inhibitors, such as VBIT-4, metformin, and VDAC1 antibodies, in our experiments suggests a therapeutic potential of targeting VDAC1 in COVID-19. These inhibitors effectively prevented cytokine-induced VDAC1 cell surface translocation, restored ATP levels, improved cell viability, and enhanced the phagocytic capacity of macrophages. Notably, metformin, a widely used drug in T2D with known effects on VDAC1 (Zhang et al. 2019), has already been associated with reduced mortality in COVID-19 patients (Bramante et al. 2021; Crouse et al. 2020). Our findings suggest that metformin’s beneficial effects may be mediated, at least in part, through its action on VDAC1, further supporting its potential as a repurposed therapy for COVID-19.

Furthermore, the inhibition of VDAC1 not only restored monocyte / macrophage cell viability and function and reduced the release of VDAC1, but also reduced the release of MCP-1, a key chemokine involved in monocyte and macrophage recruitment during inflammation. Elevated MCP-1 levels have been linked to poor outcomes in COVID-19 (Peyneau et al. 2022), and our data suggest that VDAC1 inhibition may help attenuate the hyper-inflammatory response by reducing MCP-1 release and other pro-inflammatory signals. This aligns with recent studies indicating that targeting mitochondrial pathways can modulate immune responses and improve outcomes in severe viral infections (Baik et al. 2023; Xian et al. 2022).

We acknowledge some limitations in this study. The timing and number of serum samples differed between patients. For patients who were discharged from the hospital during the study period, no blood samples were obtained until the follow-up consultation. Future studies should explore the clinical application of VDAC1 inhibitors in larger cohorts of COVID-19 patients, as well as investigate their efficacy in other inflammatory and viral diseases characterized by mitochondrial dysfunction.

Taken together, our findings provide strong support for the hypothesis that VDAC1 plays a central role in the mitochondrial dysfunction and immune dysregulation observed in severe COVID-19. The correlation between VDAC1 expression and clinical severity, along with the restoration of immune cell function by VDAC1 inhibitors, highlights the potential of VDAC1 as both a biomarker and a therapeutic target in COVID-19.

## Conclusion

In conclusion, our study underscores the importance of mitochondria’s vital role in maintaining metabolic processes and controlling cell death, particularly an altered VDAC1 expression and localization in immune cells, in the pathogenesis of severe COVID-19. The therapeutic targeting of VDAC1 would offer a promising avenue for mitigating the excessive inflammation and immune cell dysfunction that contribute to not only poor outcomes in critically ill COVID- 19 patients but also in other human diseases involving dysfunctional immunity. Further research is warranted to fully elucidate the molecular mechanisms underlying VDAC1’s role in COVID- 19 and other immune dysfunction to develop effective strategies for its clinical application.

## Acknowledgements

We would like to thank the health care workers at the Clinic of Infectious Diseases, Lund, Sweden for the assistance of blood sampling. We also thank Anna Maria Ramsay for skilled and dedicated technical assistance.

## Funding

The research was funded by the Swedish Research Council, Mats Paulsson foundation, Forget Foundation, EFSD/Lilly European Diabetes Research Programme, Novo Nordic Foundation; Swedish diabetes foundation, Bo & Kerstin hjelt Foundation for Diabetes type 2, Gunvor och Josef Anérs stiftelse and Lund University Diabetes Centre (LUDC/EXODIAB/LUDC-IRC).

## Data Availability

All relevant data are within the paper.

## Author contributions

M.S. conceived and designed the experiments. M.S., A.S., and C.B.W. supervised the project. M.S., P.D., and A.S. performed the experiments and analysed the data. K.L. was involved as an immunologist and macrophage expert. M.R. collected the clinical samples, and A.N. purified the PBMC. M.S. and A.S. wrote the first draft of the manuscript, which was substantially revised by C.B.W. All co-authors proofread the manuscript and provided valuable suggestions.

## Declaration of Interests

A.S. and C.B.W. are co-founders and shareholders of Abarceo Pharma, Malmö, Sweden. C.B.W. is employed by Abarceo Pharma and is a board member of the company.

## Notes

### Competing Interest Statement

A.S. and C.B.W. are co-founders and shareholders of Abarceo Pharma, Malmo, Sweden. C.B.W. is employed by Abarceo Pharma and is a board member of the company.

## References

1. Ajaz, S., M. J. McPhail, K. K. Singh, S. Mujib, F. M. Trovato, S. Napoli, and K. Agarwal. 2021. ’Mitochondrial metabolic manipulation by SARS-CoV-2 in peripheral blood mononuclear cells of patients with COVID-19’, Am J Physiol Cell Physiol, 320: C57–C65.

2. Akanda, N., R. Tofighi, J. Brask, C. Tamm, F. Elinder, and S. Ceccatelli. 2008. ’Voltage-dependent anion channels (VDAC) in the plasma membrane play a critical role in apoptosis in differentiated hippocampal neurons but not in neural stem cells’, Cell Cycle, 7: 3225–34.

3. Arteaga-Blanco, L. A., J. R. Temerozo, L. P. S. Tine, L. Dantas-Pereira, C. Q. Sacramento, N. Fintelman-Rodrigues, B. M. Toja, S. S. Gomes Dias, C. S. de Freitas, C. C. Espirito-Santo, Y. P. Silva, R. L. Frozza, P. T. Bozza, R. F. S. Menna-Barreto, T. M. L. Souza, and D. C. Bou-Habib. 2024. ’Extracellular vesicles from primary human macrophages stimulated with VIP or PACAP mediate anti-SARS-CoV-2 activities in monocytes through NF-kappaB signaling pathway’, Microbes Infect: 105400.

4. Baik, S. H., V. K. Ramanujan, C. Becker, S. Fett, D. M. Underhill, and A. J. Wolf. 2023. ’Hexokinase dissociation from mitochondria promotes oligomerization of VDAC that facilitates NLRP3 inflammasome assembly and activation’, Sci Immunol, 8: eade7652.

5. Ben-Hail, D., R. Begas-Shvartz, M. Shalev, A. Shteinfer-Kuzmine, A. Gruzman, S. Reina, V. De Pinto, and V. Shoshan-Barmatz. 2016. ’Novel Compounds Targeting the Mitochondrial Protein VDAC1 Inhibit Apoptosis and Protect against Mitochondrial Dysfunction’, J Biol Chem, 291: 24986–5003.

6. Bhowal, C., S. Ghosh, D. Ghatak, and R. De. 2023. ’Pathophysiological involvement of host mitochondria in SARS-CoV-2 infection that causes COVID-19: a comprehensive evidential insight’, Mol Cell Biochem, 478: 1325–43.

7. Bisen, A. C., S. Agrawal, S. N. Sanap, H. G. Ravi Kumar, N. Kumar, R. Gupta, and R. S. Bhatta. 2023. ’COVID-19 retreats and world recovers: A silver lining in the dark cloud’, Health Care Sci, 2: 264–85.

8. Bläckberg, Anna, Nils Fernström, Emma Sarbrant, Magnus Rasmussen, and Torgny Sunnerhagen. 2021. ’Antibody kinetics and clinical course of COVID-19 a prospective observational study’, PLoS One, 16: e0248918.

9. Bramante, C. T., N. E. Ingraham, T. A. Murray, S. Marmor, S. Hovertsen, J. Gronski, C. McNeil, R. Feng, G. Guzman, N. Abdelwahab, S. King, L. Tamariz, T. Meehan, K. M. Pendleton, B. Benson, D. Vojta, and C. J. Tignanelli. 2021. ’Metformin and risk of mortality in patients hospitalised with COVID-19: a retrospective cohort analysis’, Lancet Healthy Longev, 2: e34– e41.

10. Campbell, E. L., and S. P. Colgan. 2015. ’Neutrophils and inflammatory metabolism in antimicrobial functions of the mucosa’, J Leukoc Biol, 98: 517–22.

11. Chen, L., H. G. Liu, W. Liu, J. Liu, K. Liu, J. Shang, Y. Deng, and S. Wei. 2020. ’[Analysis of clinical features of 29 patients with 2019 novel coronavirus pneumonia]’, Zhonghua Jie He He Hu Xi Za Zhi, 43: 203–08.

12. Codo, A. C., G. G. Davanzo, L. B. Monteiro, G. F. de Souza, S. P. Muraro, J. V. Virgilio-da-Silva, J. S. Prodonoff, V. C. Carregari, C. A. O. de Biagi Junior, F. Crunfli, J. L. Jimenez Restrepo, P. H. Vendramini, G. Reis-de-Oliveira, K. Bispo Dos Santos, D. A. Toledo-Teixeira, P. L. Parise, M. C. Martini, R. E. Marques, H. R. Carmo, A. Borin, L. D. Coimbra, V. O. Boldrini, N. S. Brunetti, A. S. Vieira, E. Mansour, R. G. Ulaf, A. F. Bernardes, T. A. Nunes, L. C. Ribeiro, A. C. Palma, M. V. Agrela, M. L. Moretti, A. C. Sposito, F. B. Pereira, L. A. Velloso, M. A. R. Vinolo, A. Damasio, J. L. Proenca-Modena, R. F. Carvalho, M. A. Mori, D. Martins-de-Souza, H. I. Nakaya, A. S. Farias, and P. M. Moraes-Vieira. 2020. ’Elevated Glucose Levels Favor SARS-CoV-2 Infection and Monocyte Response through a HIF-1alpha/Glycolysis-Dependent Axis’, Cell Metab, 32: 437–46 e5.

13. Cory, T. J., R. S. Emmons, J. R. Yarbro, K. L. Davis, and B. D. Pence. 2021. ’Metformin Suppresses Monocyte Immunometabolic Activation by SARS-CoV-2 Spike Protein Subunit 1’, Front Immunol, 12: 733921.

14. Crouse, A. B., T. Grimes, P. Li, M. Might, F. Ovalle, and A. Shalev. 2020. ’Metformin Use Is Associated With Reduced Mortality in a Diverse Population With COVID-19 and Diabetes’, Front Endocrinol (Lausanne), 11: 600439.

15. Del Valle, D. M., S. Kim-Schulze, H. H. Huang, N. D. Beckmann, S. Nirenberg, B. Wang, Y. Lavin, T. H. Swartz, D. Madduri, A. Stock, T. U. Marron, H. Xie, M. Patel, K. Tuballes, O. Van Oekelen, A. Rahman, P. Kovatch, J. A. Aberg, E. Schadt, S. Jagannath, M. Mazumdar, A. W. Charney, A. Firpo-Betancourt, D. R. Mendu, J. Jhang, D. Reich, K. Sigel, C. Cordon-Cardo, M. Feldmann, S. Parekh, M. Merad, and S. Gnjatic. 2020. ’An inflammatory cytokine signature predicts COVID-19 severity and survival’, Nat Med, 26: 1636–43.

16. Dharra, R., A. Kumar Sharma, and S. Datta. 2023. ’Emerging aspects of cytokine storm in COVID-19: The role of proinflammatory cytokines and therapeutic prospects’, Cytokine, 169: 156287.

17. Duner, Pontus, and Albert Salehi. 2020. ’COVID-19 and possible pharmacological preventive options’, Journal of Clinical Medicine Research, 12: 758.

18. Emelyanova, L., X. Bai, Y. Yan, Z. J. Bosnjak, D. Kress, C. Warner, S. Kroboth, T. Rudic, S. Kaushik, E. Stoeckl, G. R. Ross, F. Rizvi, A. J. Tajik, and A. Jahangir. 2021. ’Biphasic effect of metformin on human cardiac energetics’, Transl Res, 229: 5–23.

19. Faraj, S. S., and P. J. Jalal. 2023. ’IL1beta, IL-6, and TNF-alpha cytokines cooperate to modulate a complicated medical condition among COVID-19 patients: case-control study’, Ann Med Surg (Lond), 85: 2291–97.

20. Ferreira, A. C., V. C. Soares, I. G. de Azevedo-Quintanilha, Sdsg Dias, N. Fintelman-Rodrigues, C. Q. Sacramento, M. Mattos, C. S. de Freitas, J. R. Temerozo, L. Teixeira, E. Damaceno Hottz, E. A. Barreto, C. R. R. Pao, L. Palhinha, M. Miranda, D. C. Bou-Habib, F. A. Bozza, P. T. Bozza, and T. M. L. Souza. 2021. ’SARS-CoV-2 engages inflammasome and pyroptosis in human primary monocytes’, Cell Death Discov, 7: 43.

21. Guan, K., Z. Zheng, T. Song, X. He, C. Xu, Y. Zhang, S. Ma, Y. Wang, Q. Xu, Y. Cao, J. Li, X. Yang, X. Ge, C. Wei, and H. Zhong. 2013. ’MAVS regulates apoptotic cell death by decreasing K48- linked ubiquitination of voltage-dependent anion channel 1’, Mol Cell Biol, 33: 3137–49.

22. Guarino, F., F. Zinghirino, L. Mela, X. G. Pappalardo, F. Ichas, V. De Pinto, and A. Messina. 2020. ’NRF-1 and HIF-1alpha contribute to modulation of human VDAC1 gene promoter during starvation and hypoxia in HeLa cells’, Biochim Biophys Acta Bioenerg, 1861: 148289.

23. Guarnieri, J. W., J. M. Dybas, H. Fazelinia, M. S. Kim, J. Frere, Y. Zhang, Y. Soto Albrecht, D. G. Murdock, A. Angelin, L. N. Singh, S. L. Weiss, S. M. Best, M. T. Lott, S. Zhang, H. Cope, V. Zaksas, A. Saravia-Butler, C. Meydan, J. Foox, C. Mozsary, Y. Bram, Y. Kidane, W. Priebe, M. R. Emmett, R. Meller, S. Demharter, V. Stentoft-Hansen, M. Salvatore, D. Galeano, F. J. Enguita, P. Grabham, N. S. Trovao, U. Singh, J. Haltom, M. T. Heise, N. J. Moorman, V. K. Baxter, E. A. Madden, S. A. Taft-Benz, E. J. Anderson, W. A. Sanders, R. J. Dickmander, S. B. Baylin, E. S. Wurtele, P. M. Moraes-Vieira, D. Taylor, C. E. Mason, J. C. Schisler, R. E. Schwartz, A. Beheshti, and D. C. Wallace. 2023. ’Core mitochondrial genes are down-regulated during SARS-CoV-2 infection of rodent and human hosts’, Sci Transl Med, 15: eabq1533.

24. Guarnieri, J. W., J. A. Haltom, Y. E. S. Albrecht, T. Lie, A. Z. Olali, G. A. Widjaja, S. S. Ranshing, A. Angelin, D. Murdock, and D. C. Wallace. 2024. ’SARS-CoV-2 mitochondrial metabolic and epigenomic reprogramming in COVID-19’, Pharmacol Res, 204: 107170.

25. Gunnarsdottir, F. B., C. Hagerling, C. Bergenfelz, M. Mehmeti, E. Kallberg, R. Allaoui, S. Mohlin, S. Pahlman, C. Larsson, K. Jirstrom, D. Bexell, and K. Leandersson. 2020. ’Inflammatory macrophage derived TNFalpha downregulates estrogen receptor alpha via FOXO3a inactivation in human breast cancer cells’, Exp Cell Res, 390: 111932.

26. Gurshaney, S., A. Morales-Alvarez, K. Ezhakunnel, A. Manalo, T. H. Huynh, J. I. Abe, N. T. Le, D. Weiskopf, A. Sette, D. S. Lupu, S. J. Gardell, and H. Nguyen. 2023. ’Metabolic dysregulation impairs lymphocyte function during severe SARS-CoV-2 infection’, Commun Biol, 6: 374.

27. Hu, H., L. Guo, J. Overholser, and X. Wang. 2022. ’Mitochondrial VDAC1: A Potential Therapeutic Target of Inflammation-Related Diseases and Clinical Opportunities’, Cells, 11.

28. Huang, C., Y. Wang, X. Li, L. Ren, J. Zhao, Y. Hu, L. Zhang, G. Fan, J. Xu, X. Gu, Z. Cheng, T. Yu, J. Xia, Y. Wei, W. Wu, X. Xie, W. Yin, H. Li, M. Liu, Y. Xiao, H. Gao, L. Guo, J. Xie, G. Wang, R. Jiang, Z. Gao, Q. Jin, J. Wang, and B. Cao. 2020. ’Clinical features of patients infected with 2019 novel coronavirus in Wuhan, China’, Lancet, 395: 497–506.

29. Huang, C., Y. Yin, P. Pan, Y. Huang, S. Chen, J. Chen, J. Wang, G. Xu, X. Tao, X. Xiao, J. Li, J. Yang, Z. Jin, B. Li, Z. Tong, W. Du, L. Liu, and Z. Liu. 2023. ’The Interaction between SARS-CoV-2 Nucleocapsid Protein and UBC9 Inhibits MAVS Ubiquitination by Enhancing Its SUMOylation’, Viruses, 15.

30. Junqueira, C., A. Crespo, S. Ranjbar, L. B. de Lacerda, M. Lewandrowski, J. Ingber, B. Parry, S. Ravid, S. Clark, M. R. Schrimpf, F. Ho, C. Beakes, J. Margolin, N. Russell, K. Kays, J. Boucau, U. Das Adhikari, S. M. Vora, V. Leger, L. Gehrke, L. A. Henderson, E. Janssen, D. Kwon, C. Sander, J. Abraham, M. B. Goldberg, H. Wu, G. Mehta, S. Bell, A. E. Goldfeld, M. R. Filbin, and J. Lieberman. 2022. ’FcgammaR-mediated SARS-CoV-2 infection of monocytes activates inflammation’, Nature, 606: 576–84.

31. Karki, R., B. R. Sharma, S. Tuladhar, E. P. Williams, L. Zalduondo, P. Samir, M. Zheng, B. Sundaram, B. Banoth, R. K. S. Malireddi, P. Schreiner, G. Neale, P. Vogel, R. Webby, C. B. Jonsson, and T. D. Kanneganti. 2021. ’Synergism of TNF-alpha and IFN-gamma Triggers Inflammatory Cell Death, Tissue Damage, and Mortality in SARS-CoV-2 Infection and Cytokine Shock Syndromes’, Cell, 184: 149–68 e17.

32. Kim, J., R. Gupta, L. P. Blanco, S. Yang, A. Shteinfer-Kuzmine, K. Wang, J. Zhu, H. E. Yoon, X. Wang, M. Kerkhofs, H. Kang, A. L. Brown, S. J. Park, X. Xu, E. Zandee van Rilland, M. K. Kim, J. I. Cohen, M. J. Kaplan, V. Shoshan-Barmatz, and J. H. Chung. 2019. ’VDAC oligomers form mitochondrial pores to release mtDNA fragments and promote lupus-like disease’, Science, 366: 1531–36.

33. Kumar, P., A. Nagarajan, and P. D. Uchil. 2018. ’Analysis of Cell Viability by the Lactate Dehydrogenase Assay’, Cold Spring Harb Protoc, 2018.

34. Li, L., Y. C. Yao, X. Q. Gu, D. Che, C. Q. Ma, Z. Y. Dai, C. Li, T. Zhou, W. B. Cai, Z. H. Yang, X. Yang, and G. Q. Gao. 2014. ’Plasminogen kringle 5 induces endothelial cell apoptosis by triggering a voltage-dependent anion channel 1 (VDAC1) positive feedback loop’, J Biol Chem, 289: 32628–38.

35. Lindeboom, R. G. H., K. B. Worlock, L. M. Dratva, M. Yoshida, D. Scobie, H. R. Wagstaffe, L. Richardson, A. Wilbrey-Clark, J. L. Barnes, L. Kretschmer, K. Polanski, J. Allen-Hyttinen, P. Mehta, D. Sumanaweera, J. M. Boccacino, W. Sungnak, R. Elmentaite, N. Huang, L. Mamanova, R. Kapuge, L. Bolt, E. Prigmore, B. Killingley, M. Kalinova, M. Mayer, A. Boyers, A. Mann, L. Swadling, M. N. J. Woodall, S. Ellis, C. M. Smith, V. H. Teixeira, S. M. Janes, R. C. Chambers, M. Haniffa, A. Catchpole, R. Heyderman, M. Noursadeghi, B. Chain, A. Mayer, K. B. Meyer, C. Chiu, M. Z. Nikolic, and S. A. Teichmann. 2024. ’Human SARS-CoV-2 challenge uncovers local and systemic response dynamics’, Nature, 631: 189–98.

36. Majtnerova, P., J. Capek, F. Petira, J. Handl, and T. Rousar. 2021. ’Quantitative spectrofluorometric assay detecting nuclear condensation and fragmentation in intact cells’, Sci Rep, 11: 11921.

37. McCarthy, M.W. 2023. ’Metformin as a potential treatment for COVID-19’, Expert Opin Pharmacother, 24: 1199–203.

38. Mohammad Al-Amily, I., M. Sjögren, P. Duner, M. Tariq, C. B. Wollheim, and A. Salehi. 2023. ’Ablation of GPR56 Causes beta-Cell Dysfunction by ATP Loss through Mistargeting of Mitochondrial VDAC1 to the Plasma Membrane’, Biomolecules, 13.

39. Neumann, S., K. Kuteykin-Teplyakov, and R. Heumann. 2024. ’Neuronal Protection by Ha-RAS-GTPase Signaling through Selective Downregulation of Plasmalemmal Voltage-Dependent Anion Channel-1’, Int J Mol Sci, 25.

40. Nomani, M., M. Varahram, P. Tabarsi, S. M. Hashemian, H. Jamaati, M. Malekmohammad, M. Ghazi, I. M. Adcock, and E. Mortaz. 2021. ’Decreased neutrophil-mediated bacterial killing in COVID- 19 patients’, Scand J Immunol, 94: e13083.

41. Parkos, C. A. 2016. ’Neutrophil-Epithelial Interactions: A Double-Edged Sword’, Am J Pathol, 186: 1404–16.

42. Peyneau, M., V. Granger, P. H. Wicky, D. Khelifi-Touhami, J. F. Timsit, F. X. Lescure, Y. Yazdanpanah, A. Tran-Dinh, P. Montravers, R. C. Monteiro, S. Chollet-Martin, M. Hurtado-Nedelec, and L. de Chaisemartin. 2022. ’Innate immune deficiencies are associated with severity and poor prognosis in patients with COVID-19’, Sci Rep, 12: 638.

43. Sefik, E., R. Qu, C. Junqueira, E. Kaffe, H. Mirza, J. Zhao, J. R. Brewer, A. Han, H. R. Steach, B. Israelow, H. N. Blackburn, S. E. Velazquez, Y. G. Chen, S. Halene, A. Iwasaki, E. Meffre, M. Nussenzweig, J. Lieberman, C. B. Wilen, Y. Kluger, and R. A. Flavell. 2022. ’Inflammasome activation in infected macrophages drives COVID-19 pathology’, Nature, 606: 585–93.

44. Shalova, I. N., J. Y. Lim, M. Chittezhath, A. S. Zinkernagel, F. Beasley, E. Hernandez-Jimenez, V. Toledano, C. Cubillos-Zapata, A. Rapisarda, J. Chen, K. Duan, H. Yang, M. Poidinger, G. Melillo, V. Nizet, F. Arnalich, E. Lopez-Collazo, and S. K. Biswas. 2015. ’Human monocytes undergo functional re-programming during sepsis mediated by hypoxia-inducible factor- 1alpha’, Immunity, 42: 484–98.

45. Shoshan-Barmatz, V., V. De Pinto, M. Zweckstetter, Z. Raviv, N. Keinan, and N. Arbel. 2010. ’VDAC, a multi-functional mitochondrial protein regulating cell life and death’, Mol Aspects Med, 31: 227–85.

46. Shoshan-Barmatz, V., A. Shteinfer-Kuzmine, and A. Verma. 2020. ’VDAC1 at the Intersection of Cell Metabolism, Apoptosis, and Diseases’, Biomolecules, 10.

47. Shteinfer-Kuzmine, A., A. Verma, R. Bornshten, E. Ben Chetrit, A. Ben-Ya’acov, H. Pahima, E. Rubin, Y. Mograbi, E. Shteyer, and V. Shoshan-Barmatz. 2024. ’Elevated serum mtDNA in COVID-19 patients is linked to SARS-CoV-2 envelope protein targeting mitochondrial VDAC1, inducing apoptosis and mtDNA release’, Apoptosis, 29: 2025–46.

48. Singh, K. K., G. Chaubey, J. Y. Chen, and P. Suravajhala. 2020. ’Decoding SARS-CoV-2 hijacking of host mitochondria in COVID-19 pathogenesis’, Am J Physiol Cell Physiol, 319: C258–C67.

49. Smilansky, A., L. Dangoor, I. Nakdimon, D. Ben-Hail, D. Mizrachi, and V. Shoshan-Barmatz. 2015. ’The Voltage-dependent Anion Channel 1 Mediates Amyloid beta Toxicity and Represents a Potential Target for Alzheimer Disease Therapy’, J Biol Chem, 290: 30670–83.

50. Smith, Robin AJ, Richard C Hartley, Helena M Cocheme, and Michael P Murphy. 2012. ’Mitochondrial pharmacology’, Trends in pharmacological sciences, 33: 341–52.

51. Tariq, M., M. Sjögren, and A. Salehi. 2024. ’Sulindac prevents increased mitochondrial VDAC1 expression and cell surface mistargeting induced by pathological conditions in retinal cells’, Biochem Biophys Res Commun, 739: 150558.

52. Thompson, E. A., K. Cascino, A. A. Ordonez, W. Zhou, A. Vaghasia, A. Hamacher-Brady, N. R. Brady, I. H. Sun, R. Wang, A. Z. Rosenberg, M. Delannoy, R. Rothman, K. Fenstermacher, L. Sauer, K. Shaw-Saliba, E. M. Bloch, A. D. Redd, A. A. R. Tobian, M. Horton, K. Smith, A. Pekosz, F. R. D’Alessio, S. Yegnasubramanian, H. Ji, A. L. Cox, and J. D. Powell. 2021. ’Metabolic programs define dysfunctional immune responses in severe COVID-19 patients’, Cell Rep, 34: 108863.

53. Tomasello, F., A. Messina, L. Lartigue, L. Schembri, C. Medina, S. Reina, D. Thoraval, M. Crouzet, F. Ichas, V. De Pinto, and F. De Giorgi. 2009. ’Outer membrane VDAC1 controls permeability transition of the inner mitochondrial membrane in cellulo during stress-induced apoptosis’, Cell Res, 19: 1363–76.

54. Verma, A., S. Pittala, B. Alhozeel, A. Shteinfer-Kuzmine, E. Ohana, R. Gupta, J. H. Chung, and V. Shoshan-Barmatz. 2022. ’The role of the mitochondrial protein VDAC1 in inflammatory bowel disease: a potential therapeutic target’, Mol Ther, 30: 726–44.

55. Vora, S. M., J. Lieberman, and H. Wu. 2021. ’Inflammasome activation at the crux of severe COVID- 19’, Nat Rev Immunol, 21: 694–703.

56. Wang, M., Y. Zhao, J. Liu, and T. Li. 2022. ’SARS-CoV-2 modulation of RIG-I-MAVS signaling: Potential mechanisms of impairment on host antiviral immunity and therapeutic approaches’, MedComm Futur Med, 1: e29.

57. Xian, H., and M. Karin. 2023. ’Oxidized mitochondrial DNA: a protective signal gone awry’, Trends Immunol, 44: 188–200.

58. Xian, H., Y. Liu, A. Rundberg Nilsson, R. Gatchalian, T. R. Crother, W. G. Tourtellotte, Y. Zhang, G. R. Aleman-Muench, G. Lewis, W. Chen, S. Kang, M. Luevanos, D. Trudler, S. A. Lipton, P. Soroosh, J. Teijaro, J. C. de la Torre, M. Arditi, M. Karin, and E. Sanchez-Lopez. 2021. ’Metformin inhibition of mitochondrial ATP and DNA synthesis abrogates NLRP3 inflammasome activation and pulmonary inflammation’, Immunity, 54: 1463–77 e11.

59. Xian, H., K. Watari, E. Sanchez-Lopez, J. Offenberger, J. Onyuru, H. Sampath, W. Ying, H. M. Hoffman, G. S. Shadel, and M. Karin. 2022. ’Oxidized DNA fragments exit mitochondria via mPTP- and VDAC-dependent channels to activate NLRP3 inflammasome and interferon signaling’, Immunity, 55: 1370–85 e8.

60. Ye, Q., B. Wang, and J. Mao. 2020. ’The pathogenesis and treatment of the ‘Cytokine Storm’ in COVID- 19’, J Infect.

61. Yeung-Luk, B. H., G. A. Narayanan, B. Ghosh, A. Wally, E. Lee, M. Mokaya, E. Wankhade, R. Zhang, B. Lee, B. Park, J. Resnick, A. Jedlicka, A. Dziedzic, M. Ramanathan, S. Biswal, A. Pekosz, and V. K. Sidhaye. 2023. ’SARS-CoV-2 infection alters mitochondrial and cytoskeletal function in human respiratory epithelial cells mediated by expression of spike protein’, mBio, 14: e0082023.

62. Zhang, E., I. Mohammed Al-Amily, S. Mohammed, C. Luan, O. Asplund, M. Ahmed, Y. Ye, D. Ben-Hail, A. Soni, N. Vishnu, P. Bompada, Y. De Marinis, L. Groop, V. Shoshan-Barmatz, E. Renstrom, C. B. Wollheim, and A. Salehi. 2019. ’Preserving Insulin Secretion in Diabetes by Inhibiting VDAC1 Overexpression and Surface Translocation in beta Cells’, Cell Metab, 29: 64–77 e6.

63. Zhang, F., J. R. Mears, L. Shakib, J. I. Beynor, S. Shanaj, I. Korsunsky, A. Nathan, Arthritis Accelerating Medicines Partnership Rheumatoid, Consortium Systemic Lupus Erythematosus, L. T. Donlin, and S. Raychaudhuri. 2020. ’IFN- gamma and TNF- alpha drive a CXCL10 + CCL2 + macrophage phenotype expanded in severe COVID-19 and other diseases with tissue inflammation’, *bioRxiv*.

64. Zhu, Z., T. Cai, L. Fan, K. Lou, X. Hua, Z. Huang, and G. Gao. 2020. ’Clinical value of immune-inflammatory parameters to assess the severity of coronavirus disease 2019’, Int J Infect Dis, 95: 332–39.

