## Supplemental file for "Shedding of mitochondrial Voltage-Dependent Anion Channel-1 (VDAC1) Reflects COVID-19 Severity and Reveals Macrophage Dysfunction"

Department of Clinical Science, Division of Islet cell physiology[1], and Division of Experimental Cardiovascular research[2], Cancer immunology, Department of Translational Medicine[3], Division of Genomics, Diabetes and Endocrinology[4], Lund University, Malmö, Sweden; Department of Clinical Sciences Lund, Faculty of Medicine, Division of Infection Medicine[5], Lund University, Lund, Sweden; Department of Cell Physiology and Metabolism, University Medical Centre, Geneva, Switzerland[6]; Abarceo Pharma, Medeon Science Park, Per Albin Hanssons v. 41, 205 12 Malmö, Sweden[7]

***Running title:*** VDAC1 in the blood of COVID-19 patients

***Key words:*** *pro-inflammatory cytokines, innate immune system, adaptive immune system, metabolite conducting channel, SARS-CoV-2 virus, human macrophages.*

**** Corresponding author:***

Albert Salehi

Department of Clinical Science

Lund University, Malmö, Sweden

### Supplementary Figures

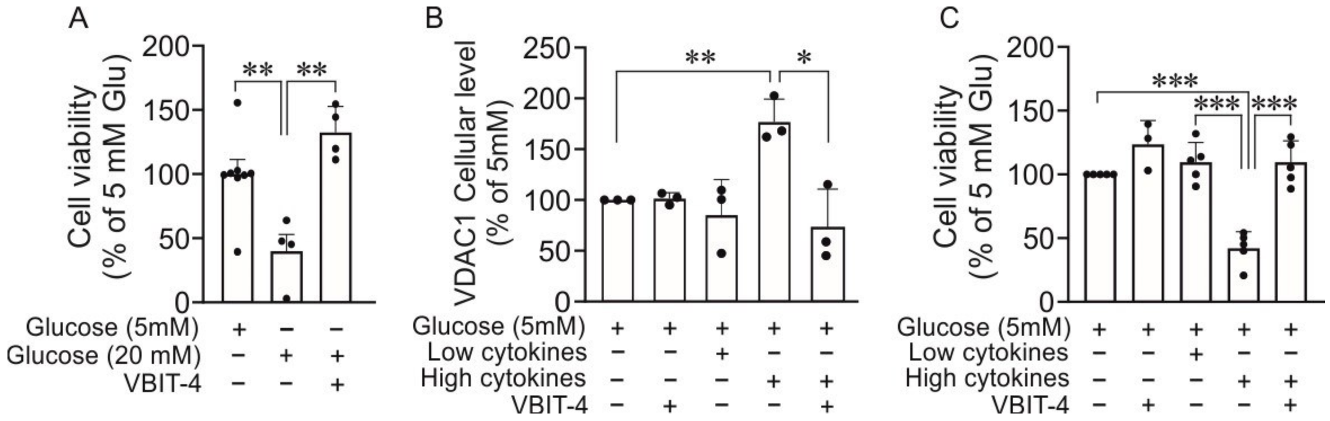


**Figure S1**. **VDAC1 expression in relation to cell viability**

Cell viability measured as reductive capacity by MTS in macrophages cultured at 5 mM or 20 mM glucose ± VBIT-4 (20 μM) for 72 h (**A**). The mean values at 5 mM glucose were kept as 100% in each experiment. Mean ± SEM from 4 independent experiments. **p < 0.01. Increased VDAC1 protein level in macrophages cultured with a high cytokine-mixture (IL-1β: 1 ng/ml, TNF-α: 250 ng/ml, INF-γ: 250 ng/ml) was suppressed by VBIT-4 (20 μM) as compared to controls (**B**). The effect of VBIT-4 alone or low cytokine concentrations (IL-1β: 0.1 ng/ml, TNF-α: 25 ng/ml, INF-γ: 25 ng/ml) on VDAC1 protein level is included. Data are mean ± SD for n=3 experiments performed at three different occasions. *p < 0.05, **p<0.01. Cell viability measured as reductive capacity by MTS in macrophages cultured at 5 mM glucose with or without VBIT-4 (20 μM), a low (IL-1β: 0.1 ng/ml, TNF-α: 25 ng/ml, INF-γ: 25 ng/ml) or a high cytokine mixture (IL-1β: 1 ng/ml, TNF-α: 250 ng/ml, INF-γ: 250 ng/ml) ±VBIT-4 for 24h (**C**). The mean values at 5 mM glucose were kept as 100% in each experiment. Data are mean ± SEM from 3-5 independent experiments. **p < 0.01, ***p<0.001.

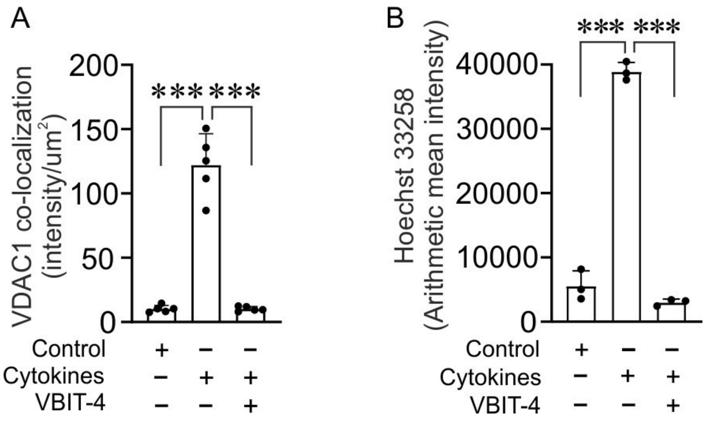


**Figure S2. Increased VDAC1 cell surface mistargeting in macrophages cultured with high concentration of cytokines**

Co-localization intensity analysis of confocal images from macrophages cultured with cytokines±VBIT-4 and double immunostained for VDAC1 and $Na^+/K^+$ ATPase (plasma membrane marker) (Shown in Figure 2) as compared to control (5 mM glucose). Mean ± SE of VDAC1/$Na^+/K^+$ ATPase co-localization intensity analysis performed on images from three independent experiments are shown. ***$p<0.001$. Detection of apoptotic nuclei exhibiting condensed or fragmented nuclei with high intense blue fluorescence (**B**). Confocal images from macrophages cultured with cytokines±VBIT-4, stained with nuclei marker (Hoechst 33258) for detection of apoptosis (Shown in Figure 2) as compared to control (5 mM glucose). Mean ± SEM of Hoechst 33258 intensity analysis performed on images from 3 independent experiments are shown. ***$p<0.001$.

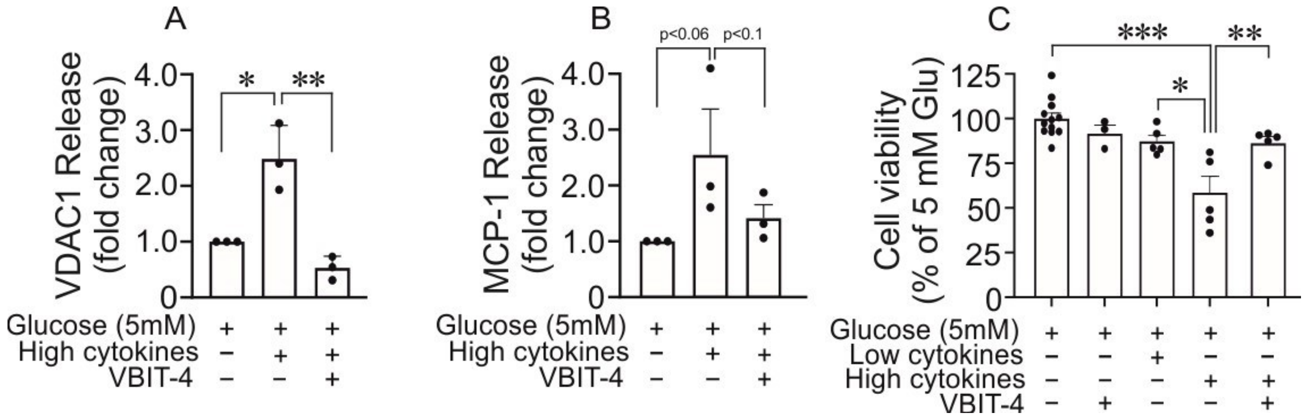


**Figure S3**. **The impact of cytokines-induced VDAC1 and MCP-1 release in monocytes in the presence or absence of VBIT-4**

Shown is the effect of cytokine mixture (IL-1β: 1 ng/ml, TNF-α: 250 ng/ml, INF-γ: 250 ng/ml) on VDAC1 (**A**) and MCP-1 (**B**) release in the presence or absence of VBIT-4 (20 µM) is compared to basal conditions (5 mM glucose) and presented as fold change. Cytokine-induced release of VDAC1 ($p<0.05$) was prevented by VBIT-4 (20 µM) ($p<0.01$) (**A**) while the cytokine-induced release MCP-1 were not significant ($p<0.063$) (**B**). Cell viability was measured as reductive capacity by MTS in monocytes cultured for 24h with a low (IL-1β: 0.1 ng/ml, TNF-α: 25 ng/ml, INF-γ: 25 ng/ml) or a high cytokine-mixture (IL-1β: 1 ng/ml, TNF-α: 250 ng/ml, INF-γ: 250 ng/ml) in the presence or absence of VBIT-4 (20 µM) (**C**). The values were normalized to 5 mM glucose control and presented as fold change. Data are mean ± SEM from 3 experiments performed at three different occasions. *$p< 0.05$, **$p <0.01$, ***$p<0.001$.
